# Whole-brain structural connectome asymmetry in autism

**DOI:** 10.1101/2023.02.15.528746

**Authors:** Seulki Yoo, Yurim Jang, Seok-Jun Hong, Hyunjin Park, Sofie L. Valk, Boris C. Bernhardt, Bo-yong Park

## Abstract

Autism spectrum disorder is a common neurodevelopmental condition that manifests as a disruption in sensory and social skills. Although it has been shown that the brain morphology of individuals with autism is asymmetric, how this differentially affects the structural connectome organization of each hemisphere remains under-investigated. We studied whole-brain structural connectivity-based brain asymmetry in 47 individuals with autism and 37 healthy controls using diffusion magnetic resonance imaging obtained from the Autism Brain Imaging Data Exchange initiative. By leveraging dimensionality reduction techniques, we constructed low-dimensional representations of structural connectivity and calculated their asymmetry index. We compared the asymmetry index between individuals with autism and neurotypical controls and found atypical structural connectome asymmetry in the sensory, default-mode, and limbic networks and the caudate in autism. Network communication provided topological underpinnings by demonstrating that the temporal and dorsolateral prefrontal regions showed reduced global network communication efficiency and decreased send-receive network navigation in the caudate region in individuals with autism. Finally, supervised machine learning revealed that structural connectome asymmetry is associated with communication-related autistic symptoms and nonverbal intelligence. Our findings provide insights into macroscale structural connectome alterations in autism and their topological underpinnings.

## Introduction

Autism spectrum disorder is a highly heritable and heterogeneous neurodevelopmental condition characterized by impaired social communication, restricted and repetitive behavior, and atypical sensory processing (Baio et al., 2018; Christensen et al., 2018; Mottron et al., 2006). These symptoms are associated with brain network disorganization and altered neuronal processing, particularly excitation/inhibition imbalances (Jou et al., 2011; Lee et al., 2017; Nelson & Valakh, 2015; Nunes et al., 2019; Sohal & Rubenstein, 2019). Previous studies have investigated the multiscale properties of the autistic brain by studying network-level brain connectomics and local microcircuit function to better understand the pathological and behavioral phenotypes of autism (Hong et al., 2019; Lee et al., 2017; Nair et al., 2013; Nelson & Valakh, 2015; Nunes et al., 2019; Park, Hong, et al., 2021).

Early studies based on magnetic resonance imaging (MRI) observed morphological and connectivity changes in the brains of individuals with autism. For example, they found increases in regional gray matter volume and cortical thickness (Khundrakpam et al., 2017; Valk et al., 2015; Zhou et al., 2014), as well as cortico-cortical functional hypoconnectivity and cortico-subcortical hyperconnectivity (Cerliani et al., 2015; Di Martino et al., 2014). Recent studies have reported abnormal brain morphology and structural network organization at the regional and large-scale network levels using a large dataset obtained from the Enhancing NeuroImaging Genetics through Meta-Analysis (ENIGMA) consortium (Postema et al., 2019; Sha et al., 2022; Van Rooij et al., 2018). On the other hand, the whole-brain connectome organization can be plotted using an advanced model based on manifold learning (*i.e*., dimensionality reduction techniques), providing low-dimensional representations of connectivity (Haak et al., 2018; Huntenburg et al., 2018; Margulies et al., 2016). Indeed, a previous study based on functional connectivity showed that the first principal component represents the hierarchical sensory-fugal axis along the cortex (Margulies et al., 2016). These techniques have been applied to study microstructural (Paquola et al., 2019; Park, Vos de Wael, et al., 2021) and structural connectivity (Park, Vos de Wael, et al., 2021), with comparable cortical axes. In particular, studies on autism datasets have used these techniques in terms of structural and functional connectivity. Functional connectome organization was atypical in low-level sensory and higher-order default-mode regions in individuals with autism compared to that in neurotypical controls (Hong et al., 2019). Moreover, alterations in the structural connectome derived from diffusion MRI tractography have been observed in sensory, somatomotor, and heteromodal association cortices (Park, Hong, et al., 2021). Likewise, individuals with autism show atypical structural and functional brain organization.

The inter-hemispheric asymmetry of brain structures, such as surface area and cortical thickness, has been recently investigated (Khundrakpam et al., 2017; Kong et al., 2018; Postema et al., 2019; Sha et al., 2022). This asymmetry provides additional information for understanding the lateralization of the brain. For example, atypical brain asymmetry patterns exists in various neuropsychiatric conditions such as schizophrenia, attention-deficit/hyperactivity disorder, and autism (Okada et al., 2016; Postema et al., 2021; Postema et al., 2019; Sha et al., 2022). Atypical asymmetry of brain structure may be associated with altered cognitive function, which may lead to disease occurrence and symptom progression. However, the asymmetry of structural connectome organization remains to be investigated. Thus, we aimed to quantitatively investigate whole-brain structural connectome asymmetry in individuals with autism using the manifold learning approach.

Network communication models may reveal the underlying topography of structural connectome asymmetry by indirectly inferring the directionality of large-scale neural signaling (Avena-Koenigsberger et al., 2018; Avena-Koenigsberger et al., 2019; Goñi et al., 2014; Seguin et al., 2018). These models measure the efficiency of information flow between different brain regions. Fast information transfer is related to the routing mechanism, whereas slow information exchange is associated with a longer path length based on the diffusion mechanism. Prior works widely adopted these network communication models to investigate the brain topology of cortical hierarchies (Vázquez-Rodríguez et al., 2019; Vézquez-Rodríguez et al., 2020) and functional dynamics (Avena-Koenigsberger et al., 2018; Park, de Wael, et al., 2021; Park, Vos de Wael, et al., 2021; Seguin et al., 2022). Thus, we hypothesized that these network communication models provide insights into the topological underpinnings of structural connectome asymmetry in autism.

Expanding upon prior studies that explored inter-hemispheric asymmetry of brain morphology (Khundrakpam et al., 2017; Kong et al., 2018; Postema et al., 2019; Sha et al., 2022), we investigated perturbations in structural connectome asymmetry in individuals with autism. We first estimated low-dimensional representations of structural connectivity based on dimensionality reduction techniques (Coifman & Lafon, 2006; Margulies et al., 2016) and assessed between-group differences in connectome asymmetry between individuals with autism and neurotypical controls. We then compared network communication measures between the groups to assess whether individuals with autism showed altered network communication. Finally, we utilized supervised machine learning to predict behavioral assessments calibrated by the Autism Diagnostic Observation Schedule (ADOS—social cognition, communication, and repeated behavior/interest sub-scores and total score) and intelligence quotient (IQ).

## Results

### Atypical structural connectome asymmetry in autism

To assess alterations in a compact feature set of structural connectivity in the low-dimensional manifold space in individuals with autism, we first estimated cortex-wide structural connectome eigenvectors by applying nonlinear dimensionality reduction techniques to a diffusion MRI tractography-derived structural connectivity matrix (Coifman & Lafon, 2006; Vos de Wael et al., 2020) (see *Methods*). The first three eigenvectors (E1, E2, and E3) explained approximately 50.1% of the total connectome information, where E1, E2, and E3 depicted anterior-posterior, superior-inferior, and lateral-medial axes, respectively, which reflect cortical hierarchy involved in cognitive processing and evolutionary adaptation (**Fig. 1A**) (Mesulam, 1998; Valk et al., 2020). Specifically, the anterior-posterior axis integrates multiple cognitive control processes from sensory stimuli to perception processing (Badre & D’esposito, 2009; Goodale & Milner, 1992), the superior-inferior axis is similar to the sensory-transmodal hierarchy model (Margulies et al., 2016), and the lateral-medial axis may describe the long-range association fibers (Tanglay et al., 2022). We then calculated the asymmetry index of the structural eigenvectors and performed multivariate analysis to compare the three asymmetry maps between individuals with autism and neurotypical controls. We observed significant between-group differences in the precuneus, rostral anterior cingulate, dorsolateral prefrontal, inferior temporal, and visual cortices (false discovery rate [FDR] < 0.05; yellow boundaries in **Fig. 1B**). By stratifying the effects according to seven intrinsic functional communities (Yeo et al., 2011) and a well-established model of large-scale cortical organization (Mesulam, 1998), we found the highest effects in the heteromodal association cortex, including the default-mode network (**Fig. 1B**).

**Fig. 1.**
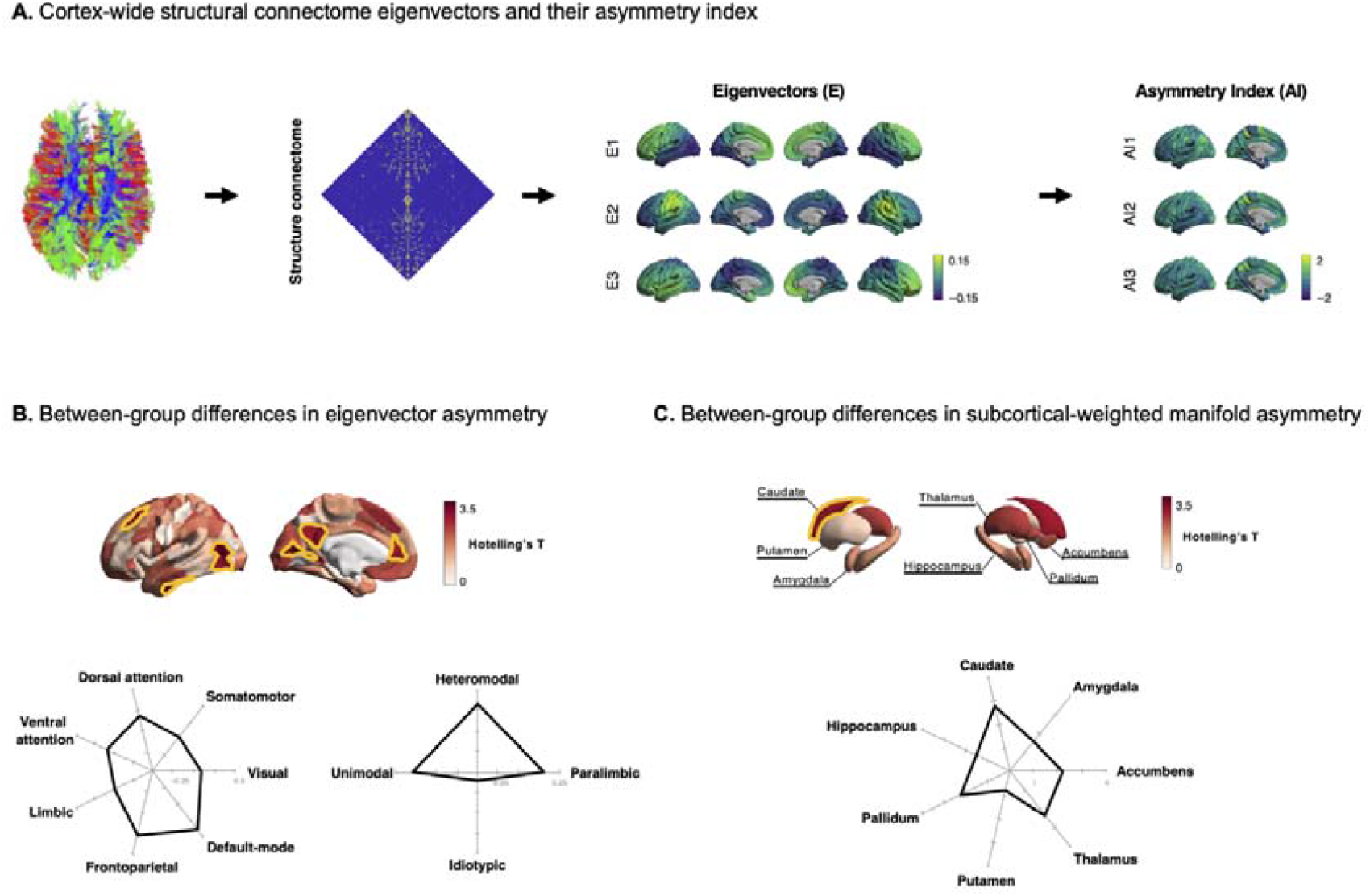
Atypical structural connectome asymmetry in individuals with autism. **(A)** Structural connectome was estimated using diffusion MRI tractography *(left)*. Three eigenvectors (E1, E2, and E3) explained approximately 50.1% of the total information shown on brain surfaces *(middle)*, and the asymmetry index was calculated *(right)*. **(B)** T-statistics of between-group differences in the asymmetry index are reported on brain surfaces, and the regions that showed significant (FDR < 0.05) effects are marked with yellow boundaries *(upper)*. We stratified the between-group difference effects of the asymmetry index according to seven intrinsic function communities and four cortical hierarchical levels *(bottom)*. **(C)** The between-group differences in the subcortical-weighted manifold asymmetry, where the significant effects are marked with a yellow boundary. *Abbreviation:* MRI, magnetic resonance imaging; FDR, false discovery rate..

In addition to cortical alterations, we investigated the between-group difference effects in subcortical areas using a subcortical-weighted manifold, reflecting subcortico-cortical connectivity weighted by the eigenvectors (Park, Hong, et al., 2021). We calculated the asymmetry of this subcortical-weighted manifold, compared individuals with autism and neurotypical controls, and identified significant effects in the caudate (FDR < 0.05; **Fig. 1C**).

### Between-group differences in structural connectivity and network communication

To assess the topological properties of structural connectome asymmetry in autism, we investigated between-group differences in structural connectivity that showed significant between-group differences in the asymmetry index (*i.e*., seed regions) and seven functional networks (Yeo et al., 2011) (*i.e*., target regions) between individuals with autism and neurotypical controls (**Fig. 2A**). We observed significant differences in connectivity between the lateral visual cortex and limbic and default-mode networks (p_perm-FDR_ = 0.007, 0.015, respectively), ventral attention network and inferior temporal (p_perm-FDR_ = 0.031) and dorsolateral prefrontal regions (p_perm-FDR_ = 0.016), and precuneus and default-mode network (p_perm-FDR_ = 0.020). Moreover, the significant difference also were seen between the caudate and visual and default-mode networks (p_perm-FDR_ = 0.044 and 0.016, respectively; **Fig. 2B**). Furthermore, we assessed network communication using the expected number of “hops” taken by a random walker to travel between two different nodes and compared both groups. We observed between-group differences in the connections between the lateral visual cortex and limbic network (p_perm-FDR_ = 0.032) and the inferior temporal cortex and four networks, including somatomotor, dorsal attention, ventral attention, and default-mode networks (p_perm-FDR_ = 0.041, 0.041, 0.046, and 0.036, respectively). Additionally, significant differences were observed between the dorsolateral prefrontal cortex somatomotor, ventral attention, frontoparietal, and default-mode networks (p_perm-FDR_ = 0.025, 0.020, 0.039, and 0.046, respectively; **Fig. 2C**). As another communication measure, we computed the navigation efficiency to compare the information transfer ability between the groups. We found significant between-group differences between the dorsolateral prefrontal cortex and dorsal attention network (p_perm-FDR_ = 0.023) and the caudate and somatomotor networks (p_perm-FDR_ = 0.022; **Fig 2D**). The findings indicate that a random walk-based multi-hop communication may be altered between the temporal and prefrontal cortices and large-scale functional brain networks. Hence, network communication efficiency decreases in individuals with autism, particularly in the temporal and prefrontal cortices.

**Fig. 2.**
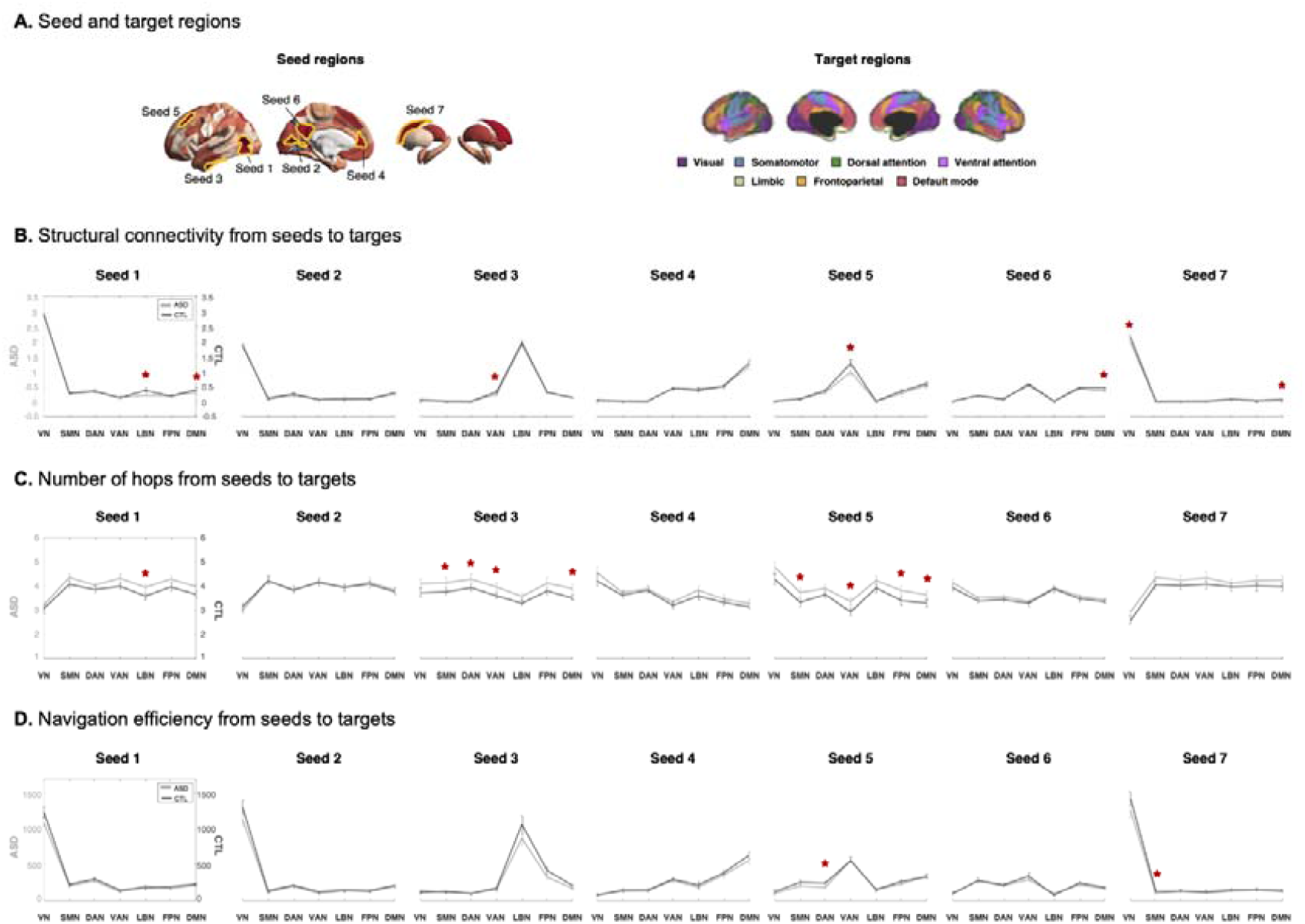
Topological underpinnings of regions showing atypical structural connectome asymmetry. **(A)** The seed and target regions are defined. **(B)** Between-group differences in seed-to-target structural connectivity, **(C)** number of hops, and **(D)** navigation efficiency between individuals with autism and neurotypical controls. Asterisks indicate significant differences (p_perm-FDR_ < 0.05). *Abbreviations:* ASD, autism spectrum disorder; CTL, control; VN, visual network; SMN, somatomotor network; DAN, dorsal attention network; VAN, ventral attention network; LBN, limbic network; FPN, frontoparietal network; DMN, default-mode network.

### Send-receive communication of atypical structural connectome asymmetry

Using navigation efficiency, we investigated the send-receive network communication of regions that showed significant between-group differences in structural connectome asymmetry (see *Methods*). We found that seed 7 (caudate) showed lower levels of both sending and receiving navigation efficiency in individuals with autism (p = 0.046 and 0.020, respectively; **Fig. 3B**). In addition, seed 1 (lateral visual cortex) showed reduced navigation efficiency (p = 0.017; **Fig. 3C**). These findings indicate that individuals with autism have altered network communication efficiency in the regions characterized by atypical structural connectome asymmetry.

**Fig. 3.**
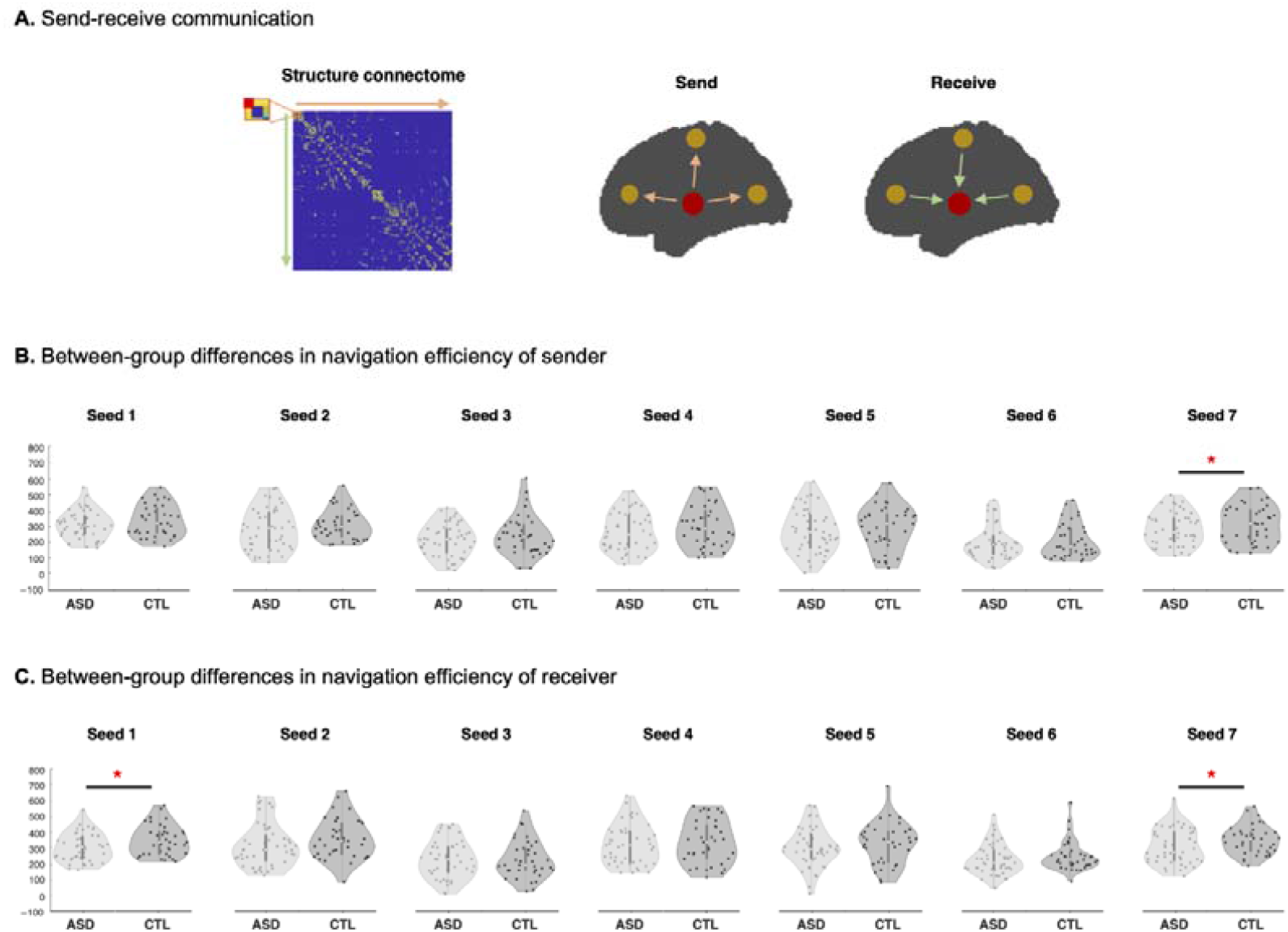
Send-receive communication of the regions showing atypical structural connectome asymmetry. **(A)** Schema of the send-receive communication of navigation efficiency. **(B)** Violin plots represent the distribution of sending and **(C)** receiving navigation efficiency of autism and control groups for each seed. *Abbreviations:* ASD, autism spectrum disorder; CTL, control.

### Behavioral phenotypes prediction

Utilizing supervised machine learning, we predicted behavioral phenotypes described by the ADOS subscales for social cognition, communication, repeated behavior/interest sub-scores, total ADOS score, and verbal and performance IQ measures and their ratio (*i.e*., verbal/performance IQ) (Hong et al., 2022) using the asymmetry index of the three eigenvectors. We significantly predicted the ADOS total score (intra-class correlation coefficient [ICC] = 0.287, p = 0.038) and communication subscore (ICC = 0.308, p = 0.028), with performance above the chance level (**Fig. 4A**). In addition, performance IQ (ICC = 0.186, p = 0.045) and IQ ratio showed significant results (ICC = 0.416, p < 0.001; **Fig. 4B**). The degree of contribution of brain regions for predicting each score was further associated with meta-analysis maps of 24 different cognitive domains derived using Neurosynth (Margulies et al., 2016), and relations to high-level sensory and perception systems were observed (**Supplementary Fig. 1**).

**Fig. 4.**
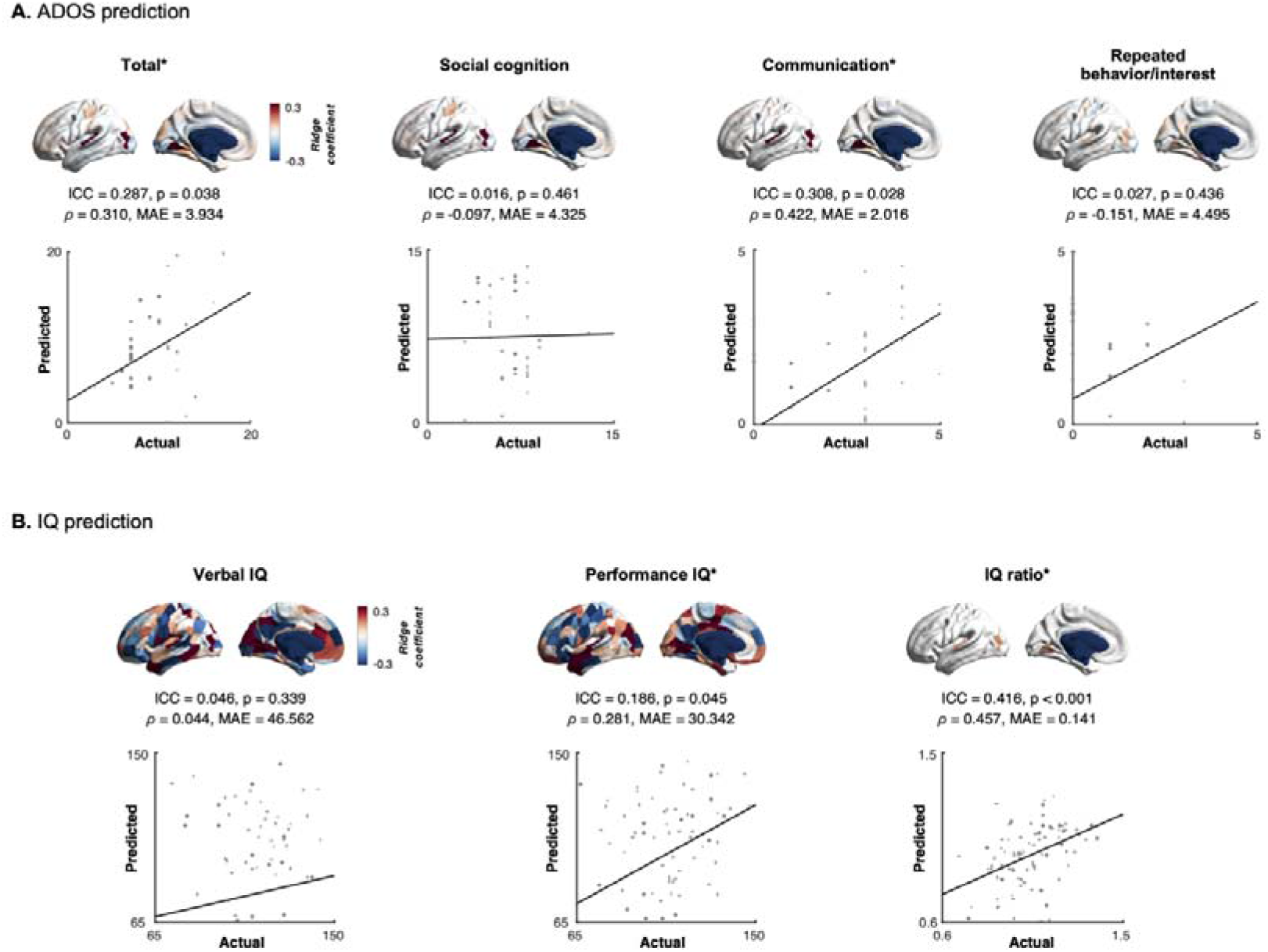
Prediction of behavioral phenotypes using structural connectome asymmetry. **(A)** We calculated linear correlations between the actual and predicted scores of ADOS total, social cognition, communication, and repeated behavior/interest scores. The coefficients of ridge regression were reported on brain surfaces. **(B)** We predicted verbal and performance IQ as well as their ratio. *Abbreviations:* ADOS, Autism Diagnostic Observation Schedule; IQ, intelligence quotient; ICC, intra-class correlation coefficient; ρ, Spearman correlation coefficient; MAE, mean absolute error.

### Sensitivity analysis

#### a) Different parcellation scales

We generated eigenvectors based on a functionally defined atlas with 100 and 300 similarly sized parcels (Schaefer et al., 2018) and assessed between-group differences in the asymmetry index of the eigenvectors between individuals with autism and neurotypical controls. Consistent results were obtained (**Supplementary Fig. 2**).

#### b) Structural parcellation scheme

We repeated the assessment of between-group differences using sub-parcellation based on the Desikan–Killiany atlas (Desikan et al., 2006; Vos de Wael et al., 2020). The findings were consistent (**Supplementary Fig. 3**), indicating the robustness of the results, regardless of the functional and structural parcellation schemes.

#### c) Head motion effect

We examined between-group differences in structural connectome asymmetry after controlling for framewise displacement (Power et al., 2012). We found consistent results, indicating that head movements do not significantly affect perturbations in interhemispheric asymmetry in individuals with autism **(Supplementary Fig. 4)**.

## Discussion

Brain network disorganization is commonly observed in individuals with autism, and asymmetry of brain morphology has been reported in multiple studies. Herein, we expanded upon prior works by systematically investigating atypical structural connectome asymmetry in individuals with autism using low-dimensional representations of structural connectivity. We observed alterations in the inter-hemispheric structural connectome asymmetry in the caudate, default-mode, and visual cortices. Network communication analyses revealed that the lateral visual, lateral temporal, and dorsolateral prefrontal cortices, as well as the caudate, showed decreases in global and local network communication efficiencies in individuals with autism. Supervised machine learning indicated that the asymmetry index may serve as a marker of autism-related communication ability and intelligence. Our findings provide insights into understanding atypical structural connectome asymmetry and network communication topology in individuals with autism.

We utilized dimensionality reduction techniques to represent cortex-wide structural connectivity with a set of macroscale eigenvectors. The selected three eigenvectors showed anterior-posterior, superior-inferior, and lateral-medial axes, consistent with the findings of previous studies (Hagmann et al., 2008; Park, Bethlehem, et al., 2021; Park, Hong, et al., 2021; Valk et al., 2020). The anterior-posterior axis is a key structure of cortical organization established in non-human primates and the human brain (Hagmann et al., 2008; Mesulam, 1998; Paquola et al., 2020). This axis consists of the rostrocaudal axis in the prefrontal cortex, which integrates multiple cognitive control processes, particularly action coupled with premotor processes (Badre & D’esposito, 2009; Braga et al., 2017; Nachev et al., 2008), and the ventral visual stream spans from the primary visual cortex to the ventral areas in the occipital and temporal cortices that implement perception processing (Borghesani et al., 2016; Goodale & Milner, 1992; Grill-Spector & Malach, 2004; Takemura et al., 2016). The superior-inferior axis resembled an established model of the sensory-transmodal hierarchy (Margulies et al., 2016), which expands from the sensorimotor area with higher myelination to heteromodal association areas with lower myelination. The lateral-medial axis is relatively under-investigated. This axis may represent long-range association fibers, such as the cingulum bundle originating from the precuneus to the orbitofrontal cortex and parahippocampal gyrus and middle longitudinal fasciculus fibers that stem from the precuneus to the lateral temporal pole, which were defined from a human *post-mortem* study (Tanglay et al., 2022).

Our findings highlighted that the principal axes of structural connectivity showed atypical asymmetry in individuals with autism, particularly in the default-mode and sensory regions. A previous study based on the ENIGMA dataset found network-level asymmetry in cortical thickness-based structural covariance in the fusiform gyrus and superior and middle frontal cortices of individuals with autism (Sha et al., 2022). Another study observed brain asymmetry in language processing-related sensory and transmodal regions in individuals with autism (Herbert et al., 2005). At the microscale, individuals with autism show less clear laminar differentiation between cortical layers IV and V (Oblak, Rosene, et al., 2011) and reduced neurotransmitter receptor binding density in the posterior cingulate cortex (Oblak, Gibbs, et al., 2011). Biophysical simulations have revealed that excitation/inhibition is related to atypical structural connectomes in autism, particularly in somatosensory and default-mode systems (Park, Hong, et al., 2021). These studies complement our findings that default-mode and sensory regions are crucial regions involved in autism at multiple scales, from macroscopic connectomics to microscale cytoarchitecture and neurotransmitters. Expanding on previous studies, we offer perspectives on atypical structural connectome asymmetry in autism.

Furthermore, we investigated the underlying connectional topology of inter-hemispheric asymmetry in autism using network communication based on a feature called network navigation, which measures the efficiency of transverse information between different brain regions (Seguin et al., 2019; Seguin et al., 2018). We assessed the global network efficiency using the number of hops between different nodes, which unfolds direct monosynaptic and indirect polysynaptic communication pathways based on random walks (Honey et al., 2009; Seguin et al., 2019; Suárez et al., 2020). A recent study based on a machine learning-based approach to building signal diffusion models suggested that a high diffusion time (*i.e*., random walk) represents polysynaptic communication pathways, whereas a low diffusion time indicates more direct monosynaptic pathways (Benkarim et al., 2022). Thus, the number of hops adopted in our study can reveal possible synaptic pathway alterations in specific brain regions in individuals with autism. Interestingly, we observed that the lateral visual, lateral temporal, and dorsolateral prefrontal cortices showed a higher number of hops toward whole-brain networks in autism, indicating a slower and inefficient signal transfer process. Studies on human and non-human primates have demonstrated that the temporal cortex is involved in higher-order sensory processing, such as language, auditory, and visual perception, and encoding of memory and emotion (Perrett et al., 1992; Perrett et al., 1984; Puce et al., 1998). Alterations in the network communication in the temporal cortex may be associated with abnormal language skills and perceptions in individuals with autism. Moreover, atypical ongoing and outgoing network communication in the dorsolateral prefrontal cortex and caudate in individuals with autism indicates susceptibility to the disease. Taken together, the inter-hemispheric structural connectome asymmetry in individuals with autism may be associated with an atypical routing network communication mechanism.

As a final analysis, we adopted supervised machine learning with cross-validation and regularization to predict the symptoms and intelligence of individuals with autism using the asymmetry index of structural connectivity. We found that asymmetry features are associated with atypical communication skills as well as performance IQ and IQ ratio, which may depend on altered cognitive and social development (Chiang et al., 2008; Wilkinson, 1998). Although more elaborate methodological approaches need to be considered to fully explain ongoing autistic symptoms and intellectual development, our findings provide insights into brain-behavior relationships in individuals with autism.

In this study, we systemically studied atypical structural connectome asymmetry in individuals with autism, investigated its topological underpinnings via network communication measures, and further stated the possibility of asymmetry as a marker for autism. Our findings may contribute to the development of brain markers for the diagnosis and prognosis of autism.

## Methods

### Study participants

We analyzed T1-weighted MRI and diffusion MRI data of 47 individuals with autism (mean ± SD age = 11.5 ± 5.7 years; 10.6 % female) and 37 healthy controls (mean ± SD age = 13 ± 4.6 years; 2.7 % female) obtained from Autism Brain Imaging Data Exchange initiative (ABIDE-II; https://fcon_1000.projects.nitrc.org/indi/abide) (Di Martino et al., 2017; Di Martino et al., 2014) (**Supplementary Table 1**). The participant list was obtained from a previous study (Park, Hong, et al., 2021), and all participants completed multimodal MRI with acceptable cortical surface reconstruction. Data were obtained from two independent sites, New York University Langone Medical Center (NYU) and Trinity College Dublin (TCD). According to gold standard diagnostic methods, all individuals with autism were diagnosed with ADOS (Lord et al., 2000) and/or Autism Diagnostic Interview-Revised (Lord et al., 1994). ABIDE data collection was performed in accordance with the local Institutional Review Board guidelines. In accordance with the HIPAA guidelines and the 1000 Functional Connectomes Project/INDI protocols, all ABIDE datasets were fully anonymized, with no protected health information included.

### Data acquisition

Multimodal MRI data from T1-weighted and diffusion MRI were acquired at two independent sites. At the NYU site, all data were scanned using a 3T Siemens Allegra scanner. The T1-weighted images were obtained using a 3D magnetization prepared rapid acquisition gradient echo (MPRAGE) sequence (repetition time (TR) = 2,530 ms; echo time (TE) = 3.25 ms; inversion time (TI) = 1,100 ms; flip angle (FA) = 7°; matrix size = 256 192; and voxel size = 1.3 1.0 1.3 mm^3^). Diffusion MRI data were obtained using a 2D spin echo-echo planar imaging (SE-EPI) sequence (TR = 5,200 ms; TE = 78 ms; matrix size = 64 64; voxel size = 3 mm^3^, isotropic; 64 directions; b-value = 1,000 s/mm^2^; and 1 b0 image). At the TCD site, the data were scanned using a 3T Philips Achieva. The T1-weighted data were acquired using a 3D MPRAGE sequence (TR = 8,400 ms; TE = 3.90 ms; TI = 1150 ms; FA = 8°; matrix = 256 256; voxel size = 0.9 mm^3^, isotropic) and diffusion MRI data using a 2D SE-EPI (TR = 20,244 ms; TE = 7.9 ms; matrix size = 124 124; voxel size = 1.94 1.94 2 mm^3^; 61 directions; b-value = 1500 s/mm^2^; and 1 b0 image).

### Data preprocessing

T1-weighted data were preprocessed using FreeSurfer. Briefly, the preprocessing steps included gradient nonuniformity correction, non-brain tissue removal, intensity normalization, and tissue segmentation. Surface reconstruction was performed using topology correction, inflation, and spherical registration to the fsaverage template space (Dale et al., 1999; Fischl et al., 2001; Fischl, Sereno, & Dale, 1999; Fischl, Sereno, Tootell, et al., 1999; Ségonne et al., 2007). The diffusion MRI data were processed using MRtrix3 (Tournier et al., 2019), which included corrections for susceptibility distortions, head motion, and eddy currents. Structural connectomes were generated from preprocessed diffusion MRI using MRtrix3 (Tournier et al., 2019). Anatomically constrained tractography was performed based on different tissue types defined using T1-weighted images (Smith et al., 2012). Multi-shell and multi-tissue response functions were estimated (Christiaens et al., 2015), and constrained spherical deconvolution and intensity normalization were performed (Jeurissen et al., 2014). The tractogram was generated based on a probabilistic approach (Tournier et al., 2019; Tournier et al., 2010; Tournier et al., 2012) with 40 million streamlines, maximum tract length of 250, and fractional anisotropy cutoff of 0.06. Subsequently, spherical deconvolution informed filtering of tractograms (SIFT2) was applied to optimize the cross-section multiplier for each streamline (Smith et al., 2015). We constructed a structural connectome by mapping the reconstructed cross-section streamlines onto the Schaefer atlas with 200 parcels (Schaefer et al., 2018) and log-transformed the values to adjust for the scale (Fornito et al., 2016) (**Fig. 1A**).

### Low-dimensional representations of structural connectivity

We estimated cortex-wide low-dimensional representations of structural connectivity (*i.e*., eigenvectors) using nonlinear dimensionality reduction techniques implemented in BrainSpace (https://github.com/MICA-MNI/BrainSpace) (Vos de Wael et al., 2020) (**Fig. 1A**). Specifically, we generated a group representative structural connectivity matrix based on distance-dependent thresholding that preserves long-range connections (Betzel et al., 2019) and estimated the eigenvectors via diffusion map embedding (Coifman & Lafon, 2006). The diffusion map embedding algorithm is robust to noise and is computationally efficient compared to other nonlinear manifold learning techniques (Tenenbaum et al., 2000; Von Luxburg, 2007). It is controlled by two parameters, α and *t*, where α controls the influence of the density of sampling points on the manifold (α = 0, maximal influence; α = 1, no influence) and *t* controls the scale of the eigenvalues of the diffusion operator. We set α = 0.5 and *t* = 0 to retain the global relations between data points in the embedded space, following prior applications (Hong et al., 2019; Margulies et al., 2016; Paquola et al., 2019; Park, Hong, et al., 2021; Vos de Wael et al., 2020). After generating the template eigenvectors, individual eigenvectors were estimated and aligned to the template via Procrustes alignment (Langs et al., 2015; Vos de Wael et al., 2020).

### Structural connectome asymmetry and between-group differences

We adopted the Schaefer atlas, which considers the asymmetric distribution of parcels (Schaefer et al., 2018), and first matched the parcels of the left and right hemispheres based on their overlap ratios. We then calculated the asymmetry index of each eigenvector as follows:

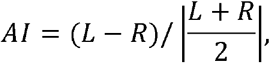

where *AI* is the asymmetry index and *L* and *R* indicate the eigenvector values of the left and right hemispheres, respectively (Bernasconi et al., 2003; Kong et al., 2018; Sarica et al., 2018). After controlling for age, sex, and site, we compared the asymmetry index between individuals with autism and neurotypical controls using multivariate analysis implemented in SurfStat (Worsley et al., 2009). Multiple comparisons across brain regions were corrected using the FDR < 0.05 (Benjamini & Hochberg, 1995). We then stratified the between-group difference effects according to seven intrinsic networks (Yeo et al., 2011) (**Fig. 1B**).

### Subcorico-cortical connectivity

In addition to the atypical asymmetry of cortico-cortical connectivity, we investigated subcortico-cortical connectivity. For each individual, subcortical regions were defined using FSL FIRST, which generates the accumbens, amygdala, pallidum, caudate, hippocampus, thalamus, and putamen (Patenaude et al., 2011). Subcortical-weighted manifolds were calculated by element-wise multiplication of subcortico-cortical connectivity with cortical eigenvectors (Park, Bethlehem, et al., 2021; Park, Hong, et al., 2021), and the nodal degree of the subcortical-weighted manifolds was calculated. We then assessed between-group differences in nodal degree values between individuals with autism and neurotypical controls after controlling for age, sex, and site (**Fig. 1C**). The FDR procedure was applied to correct for multiple comparisons across subcortical regions.

### Random walk-based network communication measures

To investigate the network communication efficiency of regions that showed atypical structural connectome asymmetry, we assessed structural connectivity and the number of hops between the seed and target regions, which measures how quickly a random walker traverses between two different nodes (Seguin et al., 2018). The seed regions showed significant between-group differences in asymmetry index between individuals with autism and neurotypical controls, and the target regions were seven intrinsic functional communities, including visual, somatomotor, dorsal attention, ventral attention, limbic, frontoparietal, and default-mode networks (Yeo et al., 2011) (**Fig. 2A**). For each participant, we stratified the structural connectivity, number of hops, and navigation efficiency between the seed and target nodes to assess the relationship between communication ability and connection strength (**Fig. 2B-D**). Statistical differences in the features between the groups were assessed using a two-sample t-test with 1,000 permutation tests. We randomly assigned autism and control groups and constructed a null distribution using t-statistics derived from the null groups. The p-value was calculated by dividing the number of permuted t-statistic values that were larger than the real t-statistic by the number of permutations. Multiple comparisons were corrected using FDR (Benjamini & Hochberg, 1995).

### Send-receive communication of navigation efficiency

To provide topological connectome underpinnings of regions that showed atypical structural connectome asymmetry, we assessed the send-receive communication of navigation efficiency (**Fig. 3A**). Navigation efficiency provides information on how to successfully find paths connecting two different nodes by capturing long and inefficient paths as well as the shortest paths (Seguin et al., 2019; Seguin et al., 2020; Seguin et al., 2018). Moreover, we can quantify how efficiently a given node sends or receives information through the observed paths (Seguin et al., 2019). Navigation efficiency was first calculated from the structural connectivity matrix for each individual using the Brain Connectivity Toolbox (https://sites.google.com/site/bctnet/) (Rubinov & Sporns, 2010). For each region that showed significant between-group differences in structural connectome asymmetry, we estimated the send and receive of navigation efficiency by calculating the row- and column-wise averages of the navigation efficiency matrix, respectively. We assessed differences in send/receive of navigation efficiency between individuals with autism and neurotypical controls using two-sample t-tests with 1,000 permutation tests.

### Prediction of behavioral phenotypes

By leveraging supervised machine learning, we predicted behavioral phenotypes of individuals with autism, including ADOS social cognition, communication, repeated behavior/interest sub-scores, total score, and verbal and performance IQ and their ratio (*i.e*., verbal/performance IQ) using the asymmetry index of the whole cortex controlled for age, sex, and site (**Fig. 4**). For each phenotype, we first assessed the regression coefficients of each independent variable using ridge regression (McDonald, 2009). The linear combination of the features and ridge coefficients was calculated to obtain the predicted phenotype score and was compared with the actual score. We performed the above process with a five-fold cross-validation, where four of the five partitions were used as training data and the remaining partition was used as test data. Performance was assessed using linear correlations between the actual and predicted phenotype scores, and significance was assessed using 1,000 permutation tests. In addition, we calculated the ICC and mean absolute error. To assess how predictive brain regions are associated with cognitive states, we associated maps of regression coefficients and 24 maps of sensory processing for cognition and memory-related processes, defined in a previous study, using Neurosynth (Margulies et al., 2016). Cognitive maps were derived from a meta-analysis to characterize the functional attributes of cognition (Yarkoni et al., 2011). The significance of the correlations was determined based on 1,000 permutation tests (**Supplementary Fig. 1**).

## Data availability

Imaging and phenotypic data were provided, in part, by the Autism Brain Imaging Data Exchange initiative (ABIDE-II; https://fcon_1000.projects.nitrc.org/indi/abide/).

## Code availability

The codes for eigenvector generation are available at https://github.com/MICA-MNI/BrainSpace, codes for calculating network communication measures are available at https://sites.google.com/site/bctnet/, and codes for statistical analyses are available at https://github.com/MICA-MNI/ENIGMA.

## Funding

Bo-yong Park received funding from the National Research Foundation of Korea (NRF-2021R1F1A1052303; NRF-2022R1A5A7033499), Institute for Information and Communications Technology Planning and Evaluation (IITP) funded by the Korea Government (MSIT) (No. 2022-0-00448, Deep Total Recall: Continual Learning for Human-Like Recall of Artificial Neural Networks; No. RS-2022-00155915, Artificial Intelligence Convergence Innovation Human Resources Development (Inha University); No. 2021-0-02068, Artificial Intelligence Innovation Hub), and Institute for Basic Science (IBS-R015-D1).

## Conflict of interest

All authors declare no conflicts of interest.

## Supplementary information

**Supplementary Table 1.** Demographic information of study participants. Means and SDs are reported.

| Information |  | NYU |  | TCD |  | p-value |
| --- | --- | --- | --- | --- | --- | --- |
| Number<br>(Autism/Control) |  | 29/18 |  | 18/19 |  | 0.232* |
| Age | Autism | 9.61 ± 6.16 | p = 0.807 | 14.46 ± 3.30 | p = 0.209 | 0.004 |
|  | Control | 10.01 ± 3.95 |  | 15.83 ± 3.21 |  | <0.001 |
| Sex<br>(male:female) | Autism | 24:5 | p = 0.243* | 18:0 | p = 1* | 0.062* |
|  | Control | 17:1 |  | 19:0 |  | 0.298* |
| ADOS — Total |  | 10.00 ± 3.36 |  | 8.72 ± 2.44 |  | 0.192 |
| ADOS — Social cognition |  | 7.50 ± 2.09 |  | 5.78 ± 2.37 |  | 0.023 |
| ADOS — Communication |  | 2.5 ± 1.70 |  | 2.94 ± 0.87 |  | 0.326 |
| ADOS — Repeated<br>behavior/interest |  | 1.40 ± 1.27 |  | 0.22 ± 0.55 |  | <0.001 |
| Verbal IQ |  | 108.55 ± 17.34 |  | 115.73 ± 14.06 |  | 0.044 |
| Performance IQ |  | 106.66 ± 19.86 |  | 113.35 ± 12.75 |  | 0.079 |
| IQ ratio<br>(verbal/performance IQ) |  | 1.03 ± 0.15 |  | 1.03 ± 0.13 |  | 0.868 |

\* Chi-squared
*Abbreviations:* SD, standard deviation; NYU, New York University Langone Medical Center; TCD, Trinity College Dublin; ADOS, Autism Diagnostic Observation Schedule; IQ, intelligence quotient.

**Supplementary Fig. 1.**
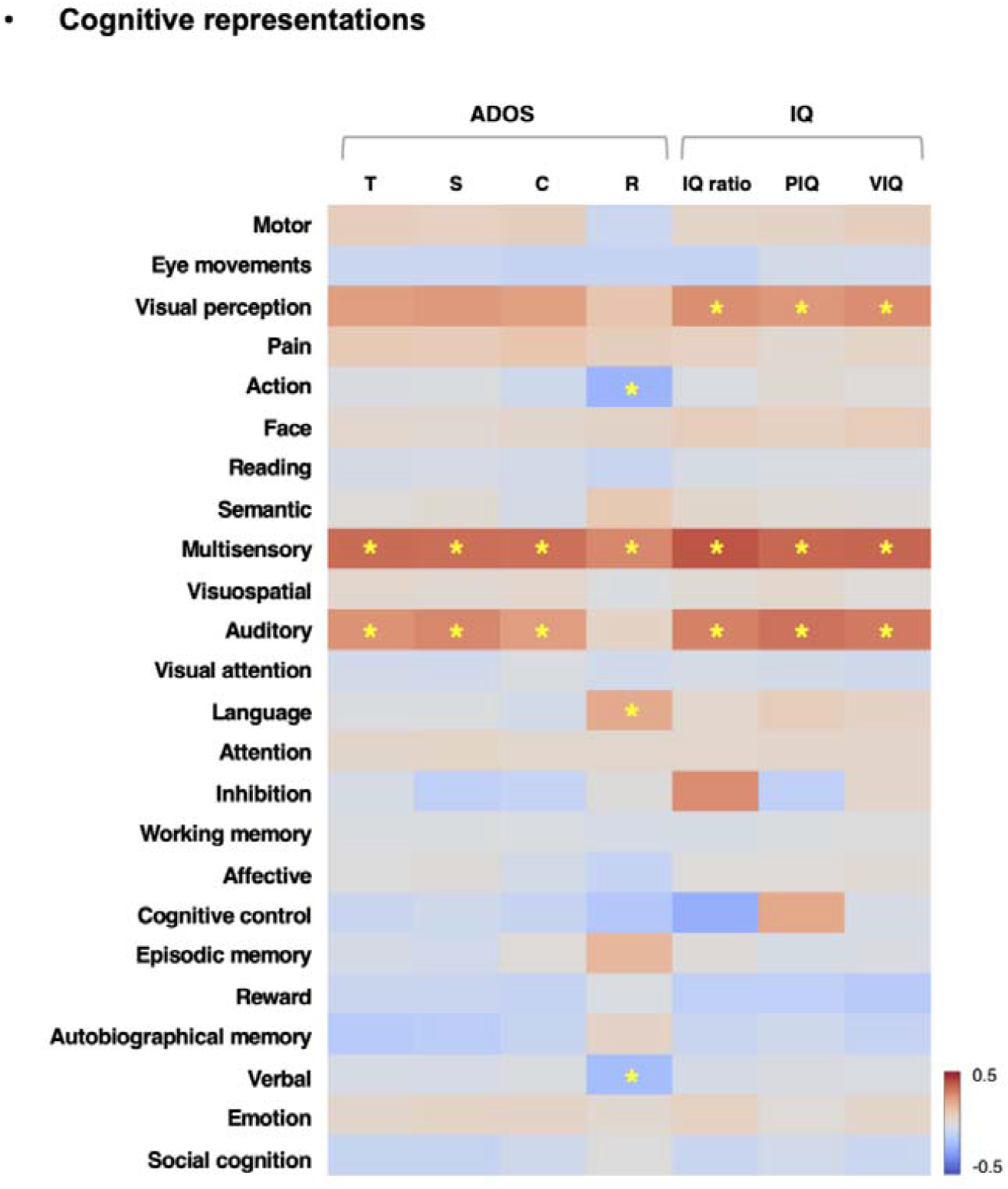
Correlation between the regression coefficients and cognitive maps derived from NeuroSynth (Yarkoni et al., 2011). The asterisks indicate p_perm_ < 0.05. *Abbreviations:* ADOS-T, Autism Diagnostic Observation Schedule total; S, social cognition; C, communication; R, repeated behavior/interest; IQ, intelligence quotient; PIQ, performance intelligence quotient; VIQ, verbal intelligence quotient.

**Supplementary Fig. 2.**
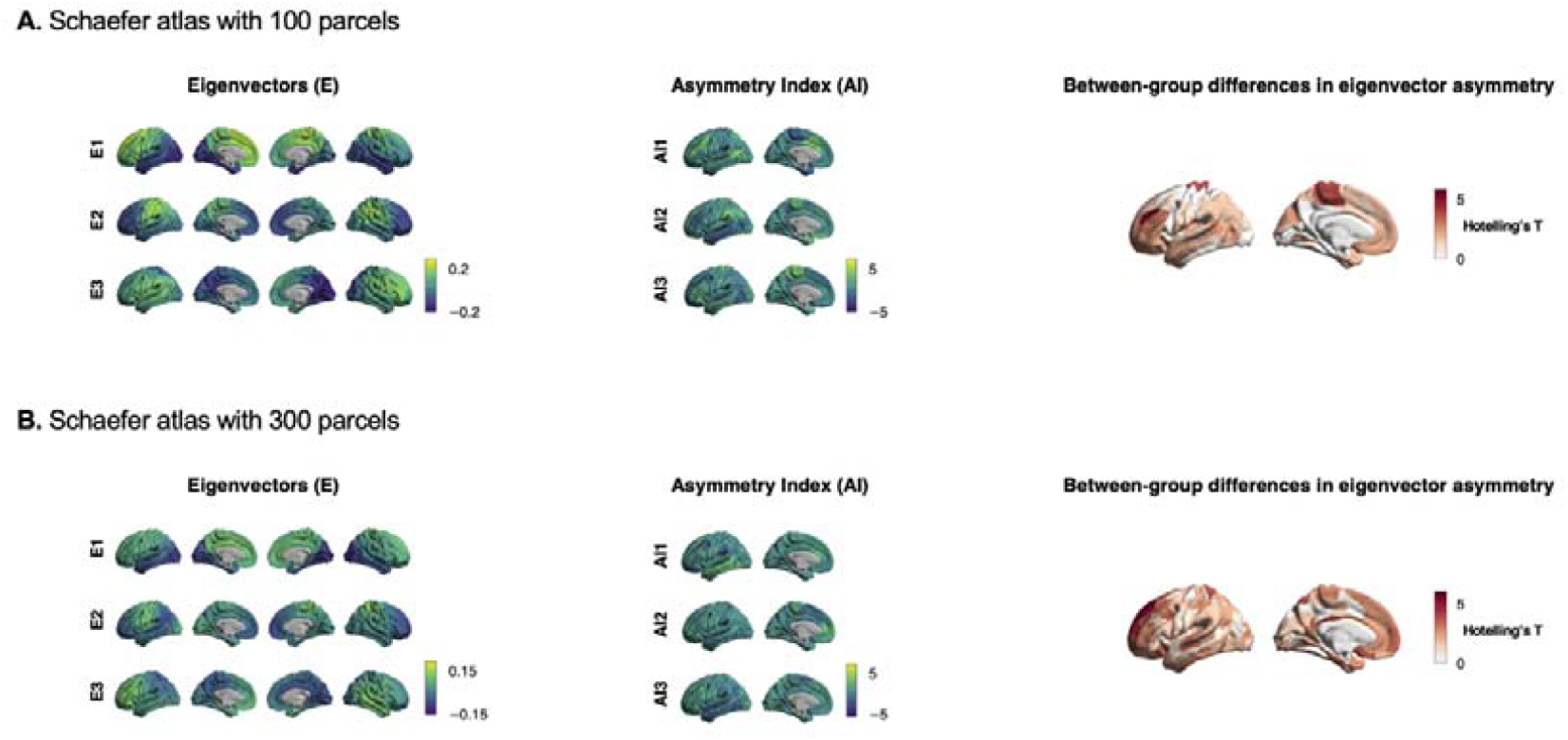
Atypical structural connectome asymmetry in individuals with autism using different spatial scales. **(A)** Three eigenvectors and their asymmetry index as well as the between-group differences in the asymmetry index between individuals with autism and neurotypical controls using Schaefer atlas with 100 and **(B)** 300 parcels (Schaefer et al., 2018).

**Supplementary Fig. 3.**
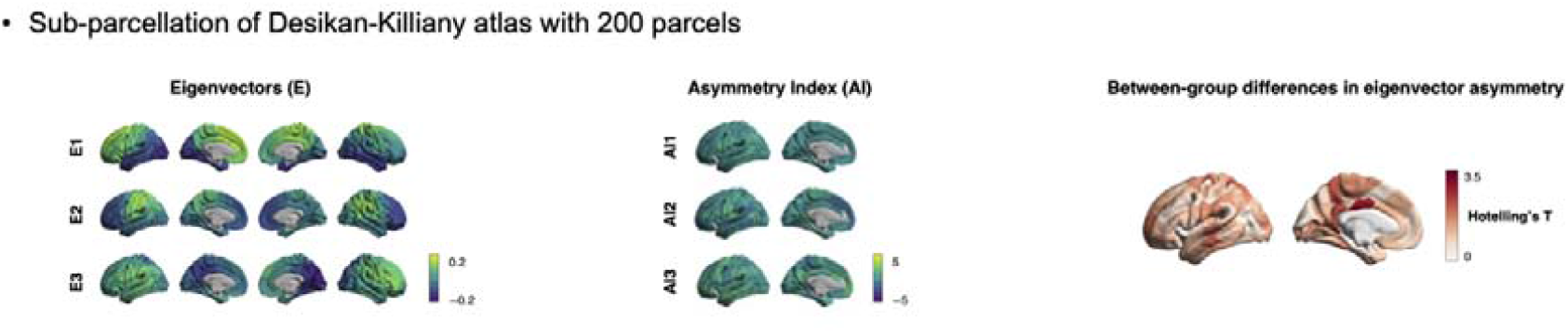
Atypical structural connectome asymmetry in individuals with autism using another parcellation scheme. A sub-parcellation scheme based on the Desikan–Killiany atlas was used (Desikan et al., 2006; Vos de Wael et al., 2020).

**Supplementary Fig. 4.**
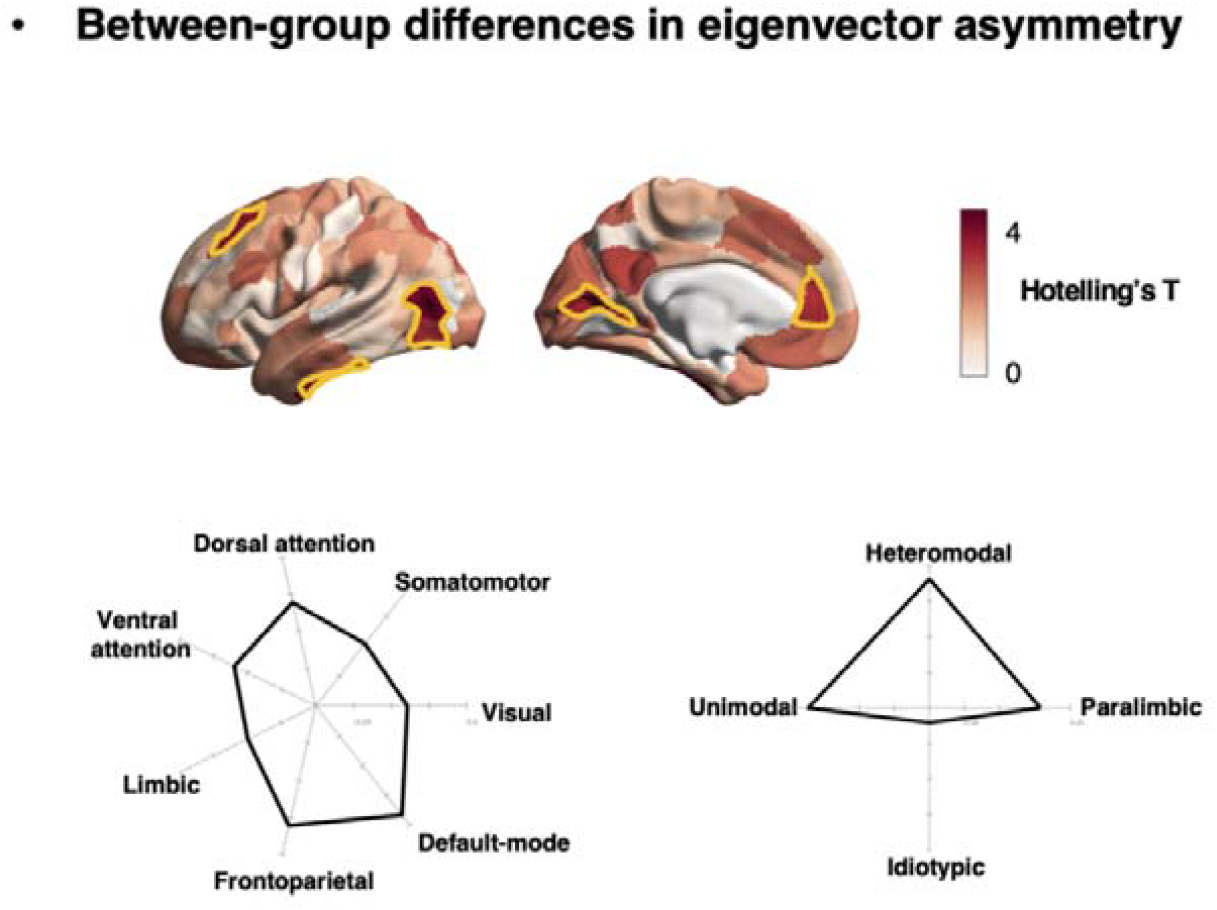
Structural connectome asymmetry controlled for head motion. The t-statistics derived from the multivariate group comparison between individuals with autism and neurotypical controls using the asymmetry index of eigenvectors after controlling for the head motion. The yellow boundaries indicate significant (false discovery rate < 0.05) between-group differences. We stratified the effects along functional communities and cortical hierarchical levels in the spider plots.

## Notes

### Competing Interest Statement

The authors have declared no competing interest.

## References

Avena-Koenigsberger, A., Misic, B., & Sporns, O. (2018). Communication dynamics in complex brain networks. Nature reviews neuroscience, 19(1), 17–33.

Avena-Koenigsberger, A., Yan, X., Kolchinsky, A., van den Heuvel, M. P., Hagmann, P., & Sporns, O. (2019). A spectrum of routing strategies for brain networks. PLoS computational biology, 15(3), e1006833.

Badre, D., & D’esposito, M. (2009). Is the rostro-caudal axis of the frontal lobe hierarchical? Nature reviews neuroscience, 10(9), 659–669.

Baio, J., Wiggins, L., Christensen, D. L., Maenner, M. J., Daniels, J., Warren, Z., Kurzius-Spencer, M., Zahorodny, W., Rosenberg, C. R., & White, T. (2018). Prevalence of autism spectrum disorder among children aged 8 years—autism and developmental disabilities monitoring network, 11 sites, United States, 2014. MMWR Surveillance Summaries, 67(6), 1.

Benjamini, Y., & Hochberg, Y. (1995). Controlling the false discovery rate: a practical and powerful approach to multiple testing. Journal of the Royal statistical society: series B (Methodological), 57(1), 289–300.

Benkarim, O., Paquola, C., Park, B.-y., Royer, J., Rodríguez-Cruces, R., de Wael, R. V., Misic, B., Piella, G., & Bernhardt, B. C. (2022). A Riemannian approach to predicting brain function from the structural connectome. Neuroimage, 119299.

Bernasconi, N., Bernasconi, A., Caramanos, Z., Antel, S., Andermann, F., & Arnold, D. L. (2003). Mesial temporal damage in temporal lobe epilepsy: a volumetric MRI study of the hippocampus, amygdala and parahippocampal region. Brain, 126(2), 462–469.

Betzel, R. F., Griffa, A., Hagmann, P., & Mišic, B. (2019). Distance-dependent consensus thresholds for generating group-representative structural brain networks. Network Neuroscience, 3(2), 475–496.

Borghesani, V., Pedregosa, F., Buiatti, M., Amadon, A., Eger, E., & Piazza, M. (2016). Word meaning in the ventral visual path: a perceptual to conceptual gradient of semantic coding. Neuroimage, 143, 128–140.

Braga, R. M., Hellyer, P. J., Wise, R. J., & Leech, R. (2017). Auditory and visual connectivity gradients in frontoparietal cortex. Human brain mapping, 38(1), 255–270.

Cerliani, L., Mennes, M., Thomas, R. M., Di Martino, A., Thioux, M., & Keysers, C. (2015). Increased functional connectivity between subcortical and cortical resting-state networks in autism spectrum disorder. JAMA psychiatry, 72(8), 767–777.

Chiang, C.-H., Soong, W.-T., Lin, T.-L., & Rogers, S. J. (2008). Nonverbal communication skills in young children with autism. Journal of autism and developmental disorders, 38(10), 1898–1906.

Christensen, D. L., Braun, K. V. N., Baio, J., Bilder, D., Charles, J., Constantino, J. N., Daniels, J., Durkin, M. S., Fitzgerald, R. T., & Kurzius-Spencer, M. (2018). Prevalence and characteristics of autism spectrum disorder among children aged 8 years—autism and developmental disabilities monitoring network, 11 sites, United States, 2012. MMWR Surveillance Summaries, 65(13), 1.

Christiaens, D., Reisert, M., Dhollander, T., Sunaert, S., Suetens, P., & Maes, F. (2015). Global tractography of multi-shell diffusion-weighted imaging data using a multi-tissue model. Neuroimage, 123, 89–101.

Coifman, R. R., & Lafon, S. (2006). Diffusion maps. Applied and computational harmonic analysis, 21(1), 5–30.

Dale, A. M., Fischl, B., & Sereno, M. I. (1999). Cortical surface-based analysis: I. Segmentation and surface reconstruction. Neuroimage, 9(2), 179–194.

Desikan, R. S., Ségonne, F., Fischl, B., Quinn, B. T., Dickerson, B. C., Blacker, D., Buckner, R. L., Dale, A. M., Maguire, R. P., & Hyman, B. T. (2006). An automated labeling system for subdividing the human cerebral cortex on MRI scans into gyral based regions of interest. Neuroimage, 31(3), 968–980.

Di Martino, A., O’connor, D., Chen, B., Alaerts, K., Anderson, J. S., Assaf, M., Balsters, J. H., Baxter, L., Beggiato, A., & Bernaerts, S. (2017). Enhancing studies of the connectome in autism using the autism brain imaging data exchange II. Scientific data, 4(1), 1–15.

Di Martino, A., Yan, C.-G., Li, Q., Denio, E., Castellanos, F. X., Alaerts, K., Anderson, J. S., Assaf, M., Bookheimer, S. Y., & Dapretto, M. (2014). The autism brain imaging data exchange: towards a large-scale evaluation of the intrinsic brain architecture in autism. Molecular psychiatry, 19(6), 659–667.

Fischl, B., Liu, A., & Dale, A. M. (2001). Automated manifold surgery: constructing geometrically accurate and topologically correct models of the human cerebral cortex. IEEE transactions on medical imaging, 20(1), 70–80.

Fischl, B., Sereno, M. I., & Dale, A. M. (1999). Cortical surface-based analysis: II: inflation, flattening, and a surface-based coordinate system. Neuroimage, 9(2), 195–207.

Fischl, B., Sereno, M. I., Tootell, R. B., & Dale, A. M. (1999). HighLresolution intersubject averaging and a coordinate system for the cortical surface. Human brain mapping, 8(4), 272–284.

Fornito, A., Zalesky, A., & Bullmore, E. (2016). Fundamentals of brain network analysis. Academic Press.

Goñi, J., Van Den Heuvel, M. P., Avena-Koenigsberger, A., Velez de Mendizabal, N., Betzel, R. F., Griffa, A., Hagmann, P., Corominas-Murtra, B., Thiran, J.-P., & Sporns, O. (2014). Resting-brain functional connectivity predicted by analytic measures of network communication. Proceedings of the National Academy of Sciences, 111(2), 833–838.

Goodale, M. A., & Milner, A. D. (1992). Separate visual pathways for perception and action. Trends in neurosciences, 15(1), 20–25.

Grill-Spector, K., & Malach, R. (2004). The human visual cortex. Annu. Rev. Neurosci., 27, 649–677.

Haak, K. V., Marquand, A. F., & Beckmann, C. F. (2018). Connectopic mapping with resting-state fMRI. Neuroimage, 170, 83–94.

Hagmann, P., Cammoun, L., Gigandet, X., Meuli, R., Honey, C. J., Wedeen, V. J., & Sporns, O. (2008). Mapping the structural core of human cerebral cortex. PLoS biology, 6(7), e159.

Herbert, M. R., Ziegler, D. A., Deutsch, C., O’Brien, L. M., Kennedy, D. N., Filipek, P., Bakardjiev, A., Hodgson, J., Takeoka, M., & Makris, N. (2005). Brain asymmetries in autism and developmental language disorder: a nested whole-brain analysis. Brain, 128(1), 213–226.

Honey, C. J., Sporns, O., Cammoun, L., Gigandet, X., Thiran, J.-P., Meuli, R., & Hagmann, P. (2009). Predicting human resting-state functional connectivity from structural connectivity. Proceedings of the National Academy of Sciences, 106(6), 2035–2040.

Hong, S.-J., Mottron, L., Park, B.-y., Benkarim, O., Valk, S. L., Paquola, C., Larivière, S., Vos de Wael, R., Degré-Pelletier, J., Soulieres, I., Ramphal, B., Margolis, A., Milham, M., Di Martino, A., & Bernhardt, B. C. (2022). A convergent structure– function substrate of cognitive imbalances in autism. Cerebral cortex. https://doi.org/10.1093/cercor/bhac156

Hong, S.-J., Vos de Wael, R., Bethlehem, R. A., Lariviere, S., Paquola, C., Valk, S. L., Milham, M. P., Di Martino, A., Margulies, D. S., & Smallwood, J. (2019). Atypical functional connectome hierarchy in autism. Nature communications, 10(1), 1–13.

Huntenburg, J. M., Bazin, P.-L., & Margulies, D. S. (2018). Large-scale gradients in human cortical organization. Trends in cognitive sciences, 22(1), 21–31.

Jeurissen, B., Tournier, J.-D., Dhollander, T., Connelly, A., & Sijbers, J. (2014). Multi-tissue constrained spherical deconvolution for improved analysis of multi-shell diffusion MRI data. Neuroimage, 103, 411–426.

Jou, R. J., Jackowski, A. P., Papademetris, X., Rajeevan, N., Staib, L. H., & Volkmar, F. R. (2011). Diffusion tensor imaging in autism spectrum disorders: preliminary evidence of abnormal neural connectivity. Australian & New Zealand Journal of Psychiatry, 45(2), 153–162.

Khundrakpam, B. S., Lewis, J. D., Kostopoulos, P., Carbonell, F., & Evans, A. C. (2017). Cortical thickness abnormalities in autism spectrum disorders through late childhood, adolescence, and adulthood: a large-scale MRI study. Cerebral cortex, 27(3), 1721–1731.

Kong, X.-Z., Mathias, S. R., Guadalupe, T., Group, E. L. W., Glahn, D. C., Franke, B., Crivello, F., Tzourio-Mazoyer, N., Fisher, S. E., & Thompson, P. M. (2018). Mapping cortical brain asymmetry in 17,141 healthy individuals worldwide via the ENIGMA Consortium. Proceedings of the National Academy of Sciences, 115(22), E5154–E5163.

Langs, G., Golland, P., & Ghosh, S. S. (2015). Predicting activation across individuals with resting-state functional connectivity based multi-atlas label fusion. International Conference on Medical Image Computing and Computer-Assisted Intervention,

Lee, E., Lee, J., & Kim, E. (2017). Excitation/inhibition imbalance in animal models of autism spectrum disorders. Biological psychiatry, 81(10), 838–847.

Lord, C., Risi, S., Lambrecht, L., Cook, E. H., Leventhal, B. L., DiLavore, P. C., Pickles, A., & Rutter, M. (2000). The Autism Diagnostic Observation Schedule—Generic: A standard measure of social and communication deficits associated with the spectrum of autism. Journal of autism and developmental disorders, 30(3), 205–223.

Lord, C., Rutter, M., & Le Couteur, A. (1994). Autism Diagnostic Interview-Revised: a revised version of a diagnostic interview for caregivers of individuals with possible pervasive developmental disorders. Journal of autism and developmental disorders, 24(5), 659–685.

Margulies, D. S., Ghosh, S. S., Goulas, A., Falkiewicz, M., Huntenburg, J. M., Langs, G., Bezgin, G., Eickhoff, S. B., Castellanos, F. X., & Petrides, M. (2016). Situating the default-mode network along a principal gradient of macroscale cortical organization. Proceedings of the National Academy of Sciences, 113(44), 12574–12579.

McDonald, G. C. (2009). Ridge regression. Wiley Interdisciplinary Reviews: Computational Statistics, 1(1), 93–100.

Mesulam, M.-M. (1998). From sensation to cognition. Brain: a journal of neurology, 121(6), 1013–1052.

Mottron, L., Dawson, M., Soulières, I., Hubert, B., & Burack, J. (2006). Enhanced perceptual functioning in autism: An update, and eight principles of autistic perception. Journal of autism and developmental disorders, 36(1), 27–43.

Nachev, P., Kennard, C., & Husain, M. (2008). Functional role of the supplementary and pre-supplementary motor areas. Nature reviews neuroscience, 9(11), 856–869.

Nair, A., Treiber, J. M., Shukla, D. K., Shih, P., & Müller, R.-A. (2013). Impaired thalamocortical connectivity in autism spectrum disorder: a study of functional and anatomical connectivity. Brain, 136(6), 1942–1955.

Nelson, S. B., & Valakh, V. (2015). Excitatory/inhibitory balance and circuit homeostasis in autism spectrum disorders. Neuron, 87(4), 684–698.

Nunes, A. S., Peatfield, N., Vakorin, V., & Doesburg, S. M. (2019). Idiosyncratic organization of cortical networks in autism spectrum disorder. Neuroimage, 190, 182–190.

Oblak, A. L., Gibbs, T. T., & Blatt, G. J. (2011). Reduced GABAA receptors and benzodiazepine binding sites in the posterior cingulate cortex and fusiform gyrus in autism. Brain research, 1380, 218–228.

Oblak, A. L., Rosene, D. L., Kemper, T. L., Bauman, M. L., & Blatt, G. J. (2011). Altered posterior cingulate cortical cyctoarchitecture, but normal density of neurons and interneurons in the posterior cingulate cortex and fusiform gyrus in autism. Autism Research, 4(3), 200–211.

Okada, N., Fukunaga, M., Yamashita, F., Koshiyama, D., Yamamori, H., Ohi, K., Yasuda, Y., Fujimoto, M., Watanabe, Y., & Yahata, N. (2016). Abnormal asymmetries in subcortical brain volume in schizophrenia. Molecular psychiatry, 21(10), 1460–1466.

Paquola, C., Seidlitz, J., Benkarim, O., Royer, J., Klimes, P., Bethlehem, R. A., Larivière, S., Vos de Wael, R., Rodríguez-Cruces, R., & Hall, J. A. (2020). A multi-scale cortical wiring space links cellular architecture and functional dynamics in the human brain. PLoS biology, 18(11), e3000979.

Paquola, C., Vos De Wael, R., Wagstyl, K., Bethlehem, R. A., Hong, S.-J., Seidlitz, J., Bullmore, E. T., Evans, A. C., Misic, B., & Margulies, D. S. (2019). Microstructural and functional gradients are increasingly dissociated in transmodal cortices. PLoS biology, 17(5), e3000284.

Park, B.-y., Bethlehem, R. A., Paquola, C., Larivière, S., Rodríguez-Cruces, R., de Wael, R. V., Bullmore, E. T., & Bernhardt, B. C. (2021). An expanding manifold in transmodal regions characterizes adolescent reconfiguration of structural connectome organization. elife, 10.

Park, B.-y., de Wael, R. V., Paquola, C., Larivière, S., Benkarim, O., Royer, J., Tavakol, S., Cruces, R. R., Li, Q., & Valk, S. L. (2021). Signal diffusion along connectome gradients and inter-hub routing differentially contribute to dynamic human brain function. Neuroimage, 224, 117429.

Park, B.-y., Hong, S.-J., Valk, S. L., Paquola, C., Benkarim, O., Bethlehem, R. A., Di Martino, A., Milham, M. P., Gozzi, A., & Yeo, B. (2021). Differences in subcortico-cortical interactions identified from connectome and microcircuit models in autism. Nature communications, 12(1), 1–15.

Park, B.-y., Vos de Wael, R., Paquola, C., Larivière, S., Benkarim, O., Royer, J., Tavakol, S., Cruces, R. R., Li, Q., & Valk, S. L. (2021). Signal diffusion along connectome gradients and inter-hub routing differentially contribute to dynamic human brain function. Neuroimage, 224, 117429.

Patenaude, B., Smith, S. M., Kennedy, D. N., & Jenkinson, M. (2011). A Bayesian model of shape and appearance for subcortical brain segmentation. Neuroimage, 56(3), 907–922.

Perrett, D. I., Hietanen, J. K., Oram, M. W., & Benson, P. J. (1992). Organization and functions of cells responsive to faces in the temporal cortex. Philosophical transactions of the royal society of London. Series B: Biological sciences, 335(1273), 23–30.

Perrett, D. I., Smith, P., Potter, D. D., Mistlin, A., Head, A., Milner, A., & Jeeves, M. (1984). Neurones responsive to faces in the temporal cortex: studies of functional organization, sensitivity to identity and relation to perception. Human neurobiology, 3(4), 197–208.

Postema, M. C., Hoogman, M., Ambrosino, S., Asherson, P., Banaschewski, T., Bandeira, C. E., Baranov, A., Bau, C. H., Baumeister, S., & BaurLStreubel, R. (2021). Analysis of structural brain asymmetries in attentionLdeficit/hyperactivity disorder in 39 datasets. Journal of Child Psychology and Psychiatry, 62(10), 1202–1219.

Postema, M. C., Van Rooij, D., Anagnostou, E., Arango, C., Auzias, G., Behrmann, M., Calderoni, S., Calvo, R., Daly, E., & Deruelle, C. (2019). Altered structural brain asymmetry in autism spectrum disorder in a study of 54 datasets. Nature communications, 10(1), 1–12.

Power, J. D., Barnes, K. A., Snyder, A. Z., Schlaggar, B. L., & Petersen, S. E. (2012). Spurious but systematic correlations in functional connectivity MRI networks arise from subject motion. Neuroimage, 59(3), 2142–2154.

Puce, A., Allison, T., Bentin, S., Gore, J. C., & McCarthy, G. (1998). Temporal cortex activation in humans viewing eye and mouth movements. Journal of neuroscience, 18(6), 2188–2199.

Rubinov, M., & Sporns, O. (2010). Complex network measures of brain connectivity: uses and interpretations. Neuroimage, 52(3), 1059–1069.

Sarica, A., Vasta, R., Novellino, F., Vaccaro, M. G., Cerasa, A., Quattrone, A., & Initiative, A. s. D. N. (2018). MRI asymmetry index of hippocampal subfields increases through the continuum from the mild cognitive impairment to the Alzheimer’s disease. Frontiers in Neuroscience, 12, 576.

Schaefer, A., Kong, R., Gordon, E. M., Laumann, T. O., Zuo, X.-N., Holmes, A. J., Eickhoff, S. B., & Yeo, B. T. (2018). Local-global parcellation of the human cerebral cortex from intrinsic functional connectivity MRI. Cerebral cortex, 28(9), 3095–3114.

Ségonne, F., Pacheco, J., & Fischl, B. (2007). Geometrically accurate topology-correction of cortical surfaces using nonseparating loops. IEEE transactions on medical imaging, 26(4), 518–529.

Seguin, C., Razi, A., & Zalesky, A. (2019). Inferring neural signalling directionality from undirected structural connectomes. Nature communications, 10(1), 1–13.

Seguin, C., Sporns, O., Zalesky, A., & Calamante, F. (2022). Network communication models narrow the gap between the modular organization of structural and functional brain networks. Neuroimage, 257, 119323.

Seguin, C., Tian, Y., & Zalesky, A. (2020). Network communication models improve the behavioral and functional predictive utility of the human structural connectome. Network Neuroscience, 4(4), 980–1006.

Seguin, C., Van Den Heuvel, M. P., & Zalesky, A. (2018). Navigation of brain networks. Proceedings of the National Academy of Sciences, 115(24), 6297–6302.

Sha, Z., Van Rooij, D., Anagnostou, E., Arango, C., Auzias, G., Behrmann, M., Bernhardt, B., Bolte, S., Busatto, G. F., & Calderoni, S. (2022). Subtly altered topological asymmetry of brain structural covariance networks in autism spectrum disorder across 43 datasets from the ENIGMA consortium. Molecular psychiatry, 27(4), 2114–2125.

Smith, R. E., Tournier, J.-D., Calamante, F., & Connelly, A. (2012). Anatomically-constrained tractography: improved diffusion MRI streamlines tractography through effective use of anatomical information. Neuroimage, 62(3), 1924–1938.

Smith, R. E., Tournier, J.-D., Calamante, F., & Connelly, A. (2015). SIFT2: Enabling dense quantitative assessment of brain white matter connectivity using streamlines tractography. Neuroimage, 119, 338–351.

Sohal, V. S., & Rubenstein, J. L. (2019). Excitation-inhibition balance as a framework for investigating mechanisms in neuropsychiatric disorders. Molecular psychiatry, 24(9), 1248–1257.

Suárez, L. E., Markello, R. D., Betzel, R. F., & Misic, B. (2020). Linking structure and function in macroscale brain networks. Trends in cognitive sciences, 24(4), 302–315.

Takemura, H., Rokem, A., Winawer, J., Yeatman, J. D., Wandell, B. A., & Pestilli, F. (2016). A major human white matter pathway between dorsal and ventral visual cortex. Cerebral cortex, 26(5), 2205–2214.

Tanglay, O., Young, I. M., Dadario, N. B., Briggs, R. G., Fonseka, R. D., Dhanaraj, V., Hormovas, J., Lin, Y.-H., & Sughrue, M. E. (2022). Anatomy and white-matter connections of the precuneus. Brain Imaging and Behavior, 16(2), 574–586.

Tenenbaum, J. B., Silva, V. d., & Langford, J. C. (2000). A global geometric framework for nonlinear dimensionality reduction. Science, 290(5500), 2319–2323.

Tournier, J.-D., Smith, R., Raffelt, D., Tabbara, R., Dhollander, T., Pietsch, M., Christiaens, D., Jeurissen, B., Yeh, C.-H., & Connelly, A. (2019). MRtrix3: A fast, flexible and open software framework for medical image processing and visualisation. Neuroimage, 202, 116137.

Tournier, J. D., Calamante, F., & Connelly, A. (2010). Improved probabilistic streamlines tractography by 2nd order integration over fibre orientation distributions. Proceedings of the international society for magnetic resonance in medicine,

Tournier, J. D., Calamante, F., & Connelly, A. (2012). MRtrix: diffusion tractography in crossing fiber regions. International journal of imaging systems and technology, 22(1), 53–66.

Valk, S. L., Di Martino, A., Milham, M. P., & Bernhardt, B. C. (2015). Multicenter mapping of structural network alterations in autism. Human brain mapping, 36(6), 2364–2373.

Valk, S. L., Xu, T., Margulies, D. S., Masouleh, S. K., Paquola, C., Goulas, A., Kochunov, P., Smallwood, J., Yeo, B. T., & Bernhardt, B. C. (2020). Shaping brain structure: Genetic and phylogenetic axes of macroscale organization of cortical thickness. Science Advances, 6(39), eabb3417.

Van Rooij, D., Anagnostou, E., Arango, C., Auzias, G., Behrmann, M., Busatto, G. F., Calderoni, S., Daly, E., Deruelle, C., & Di Martino, A. (2018). Cortical and subcortical brain morphometry differences between patients with autism spectrum disorder and healthy individuals across the lifespan: results from the ENIGMA ASD Working Group. American Journal of Psychiatry, 175(4), 359–369.

Vázquez-Rodríguez, B., Suárez, L. E., Markello, R. D., Shafiei, G., Paquola, C., Hagmann, P., Van Den Heuvel, M. P., Bernhardt, B. C., Spreng, R. N., & Misic, B. (2019). Gradients of structure–function tethering across neocortex. Proceedings of the National Academy of Sciences, 116(42), 21219–21227.

Vézquez-Rodríguez, B., Liu, Z.-Q., Hagmann, P., & Misic, B. (2020). Signal propagation via cortical hierarchies. Network neuroscience, 4(4), 1072–1090.

Von Luxburg, U. (2007). A tutorial on spectral clustering. Statistics and computing, 17(4), 395–416.

Vos de Wael, R., Benkarim, O., Paquola, C., Lariviere, S., Royer, J., Tavakol, S., Xu, T., Hong, S.-J., Langs, G., & Valk, S. (2020). BrainSpace: a toolbox for the analysis of macroscale gradients in neuroimaging and connectomics datasets. Communications biology, 3(1), 1–10.

Wilkinson, K. M. (1998). Profiles of language and communication skills in autism. Mental retardation and developmental disabilities research reviews, 4(2), 73–79.

Worsley, K. J., Taylor, J., Carbonell, F., Chung, M., Duerden, E., Bernhardt, B., Lyttelton, O., Boucher, M., & Evans, A. (2009). A Matlab toolbox for the statistical analysis of univariate and multivariate surface and volumetric data using linear mixed effects models and random field theory. NeuroImage Organisation for Human Brain Mapping 2009 Annual Meeting,

Yarkoni, T., Poldrack, R. A., Nichols, T. E., Van Essen, D. C., & Wager, T. D. (2011). Large-scale automated synthesis of human functional neuroimaging data. Nature methods, 8(8), 665–670.

Yeo, B. T., Krienen, F. M., Sepulcre, J., Sabuncu, M. R., Lashkari, D., Hollinshead, M., Roffman, J. L., Smoller, J. W., Zöllei, L., & Polimeni, J. R. (2011). The organization of the human cerebral cortex estimated by intrinsic functional connectivity. Journal of neurophysiology.

Zhou, Y., Yu, F., & Duong, T. (2014). Multiparametric MRI characterization and prediction in autism spectrum disorder using graph theory and machine learning. PloS one, 9(6), e90405.

